# msCentipede: Modeling heterogeneity across genomic sites improves accuracy in the inference of transcription factor binding

**DOI:** 10.1101/012013

**Authors:** Anil Raj, Heejung Shim, Yoav Gilad, Jonathan K. Pritchard, Matthew Stephens

**Affiliations:** Department of Genetics, Stanford University, Stanford, CA, 94305; Department of Human Genetics, University of Chicago, Chicago, IL, 60637; Department Biology, Stanford University, Stanford, CA, 94305; Howard Hughes Medical Institute; Department of Statistics, University of Chicago, Chicago, IL, 60637

## Abstract

**Motivation:** Understanding global gene regulation depends critically on accurate annotation of regulatory elements that are functional in a given cell type. CENTIPEDE, a powerful, probabilistic framework for identifying transcription factor binding sites from tissue-specific DNase I cleavage patterns and genomic sequence content, leverages the hypersensitivity of factor-bound chromatin and the information in the DNase I spatial cleavage profile characteristic of each DNA binding protein to accurately infer functional factor binding sites. However, the model for the spatial profile in this framework underestimates the substantial variation in the DNase I cleavage profiles across factor-bound genomic locations and across replicate measurements of chromatin accessibility.

**Results:** In this work, we adapt a multi-scale modeling framework for inhomogeneous Poisson processes to better model the underlying variation in DNase I cleavage patterns across genomic locations bound by a transcription factor. In addition to modeling variation, we also model spatial structure in the heterogeneity in DNase I cleavage patterns for each factor. Using DNase-seq measurements assayed in a lymphoblastoid cell line, we demonstrate the improved performance of this model for several transcription factors by comparing against the Chip-Seq peaks for those factors. Finally, we propose an extension to this framework that allows for a more flexible background model and evaluate the additional gain in accuracy achieved when the background model parameters are estimated using DNase-seq data from naked DNA. The proposed model can also be applied to paired-end ATAC-seq and DNase-seq data in a straightforward manner.

**Availability:** msCentipede, a Python implementation of an algorithm to infer transcription factor binding using this model, is made available at https://github.com/rajanil/msCentipede

## 1 Introduction

A central challenge in modern genomics is the accurate identification of all the regulatory sequences that are active in a given cell type and a description of the mechanisms by which they regulate gene expression. One key mechanism is by recruiting transcription factors which bind to the DNA at characteristic nucleotide sequences. Chromatin immunoprecipitation followed by sequencing (ChIP-seq) provides a direct measurement of DNA sequences bound by transcription factors (either directly or through a co-factor); however, each ChIP-seq experiment provides information for only one transcription factor at a time. DNase-seq (Boyle *et al.,* 2008; Hesselberth *et al.,* 2009) provides an indirect measurement of active regulatory sequences by exploiting the increased sensitivity of nucleosome-depleted chromatin to DNase I enzyme. While DNase-seq provides information on the active regulatory regions in the genome, identifying which transcription factors are bound to these regions and their organization requires statistical modelling of the spatial structure in DNase sensitivity in active regulatory regions (Pique-Regi *et al.* 2010; Boyle *et al.,* 2011; Luo and Hartemink, 2013; Piper *et al.,* 2013; Sherwood *et al.,* 2014).

Pique-Regi *et al.* (2010) put forward a probabilistic framework to infer sequence motif instances that are bound by transcription factors by combining sequence information with the information in DNase I cleavage patterns measured from DNase-seq assays. The model, CENTIPEDE, relies on two observations: (1) chromatin around motif instances bound by transcription factors typically has higher DNase I sensitivity than chromatin around unbound motif instances, and (2) each transcription factor has a characteristic DNase I cleavage profile around bound motif instances. Based on these observations, given a putative bound motif instance, CENTIPEDE models the number of reads mapped to each base pair along a window around the motif site as a mixture of two components (bound vs unbound), and infers the probability that each site is bound. Specifically, conditional on being bound (or unbound), CENTIPEDE models (1) the total number of DNase-seq reads using a negative binomial distribution, and (2) DNase-seq read counts along a window, conditional on the total number of reads, using a multinomial distribution, with independent sets of parameters for bound and unbound sites.

The multinomial model, however, effectively assumes that given enough number of reads, the DNase I read count profiles would be the same at all bound sites. Instead, we observe that the read count profiles often have larger variation across factor-bound genomic locations than a multinomial distribution can model. Based on this, we hypothesized that improved modeling of the excess variation would likely lead to improved performance in predicting transcription factor binding.

To illustrate the excess variation, we make use of the connection between the multinomial distribution and binomial distribution. Specifically, if the read counts per base pair are multinomially distributed conditional on the total read count in a genomic window, then the number of reads mapping to the left half of the window conditional on total read count should be binomially distributed. Based on this, we can compare the true distribution of the proportion of reads mapping to the left half of a genomic window to a distribution of proportions computed by sampling read counts from a binomial distribution (see Supplementary Methods for details). Ideally, we would like to illustrate overdispersion in the read counts mapping to the left half of the window across genomic sites. However, since the total read counts vary substantially across genomic sites, we resorted to using proportions rather than absolute read counts to illustrate the overdispersion across genomic sites.

In Figure 1, we observe that the distribution of ‘true’ proportions has a higher variance than that of ‘simulated’ proportions, suggesting that the multinomial distribution is insufficient to model the variation in read profiles across genomic sites bound by the transcription factor. We also observe overdispersion in the ‘true’ proportions across multiple scales and window sizes. Furthermore, when multiple replicate DNase-seq measurements are available for the same cell type, CENTIPEDE has often been applied after pooling replicates. If there is substantial heterogeneity between replicates, then pooling replicates tends to introduce more variation in the read count profiles, exacerbating the limitation of the multinomial model in this framework.

**Figure 1:**
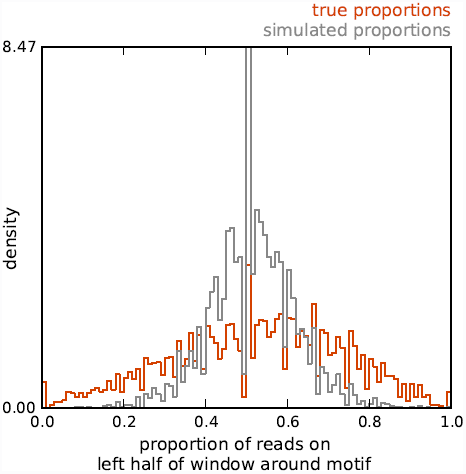
Excess variation in DNase I cleavage rate along the genome that is not explained by binomial sampling. For a set of 1000 SP1 motif instances with high ChIP-seq signal, we computed, for a 100bp window around each motif instance, the ratio of number of DNase I cuts mapped to the left half of the window to the number of DNase I cuts mapped to the entire window. The histogram of these ‘true ratios’ is shown in orange. For each window, we also simulated read counts mapping to the left half of the window by sampling from a binomial distribution whose parameter is estimated from these data (see Supplementary Methods for more details). A histogram of ‘simulated ratios’ computed from these simulated read counts is shown in gray.

Motivated by recent work applying multi-scale methods to analyses of high-throughput sequencing data (Shim and Stephens, 2013; Shim *et al.,* 2014), we use hierarchical multi-scale models to better model heterogeneity in the read profiles across genomic locations and across replicate measurements of chromatin accessibility. Modeling the data at multiple scales explicitly allows us to infer different amounts of genomewide variation at each scale, and enables the automatic identification of relevant scales during inference. In addition, the proposed multi-scale modeling framework better models spatial structure in the heterogeneity in DNase I cleavage patterns induced by the binding of a particular transcription factor. Finally, when DNase-seq data from naked DNA are available, we propose a flexible DNase I cleavage background model, the parameters of which are estimated from these additional DNase-seq data.

## 2 Models and Methods

Consider a genomic window (site) of length *L* centered around each of *N* putative binding motifs, with *L* assumed to be a power of 2 (*L* = 2^*J*^). Let 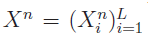 be the sequence of read counts for the *n*^th^ site, where 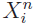 is read count at *i*^th^ base pair in the site. Let *Z*^*n*^ denote a binary indicator for whether the *n*^th^ site is bound (*Z*^*n*^ = 1). Following the model in CENTIPEDE, a mixture model at the *n*^th^ site can be written as

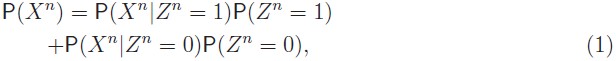

where the mixing proportion P (*Z*^*n*^ = 1) = ζ can be modeled as a logistic function of genomic information (e.g. motif position weight matrix score and motif sequence conservation score) as in CENTIPEDE.

### 2.1 msCentipede model at bound motifs

We modeled the profile of read counts at the *n*^th^ site *X*^*n*^ conditional on *Z*^*n*^ = 1 using a Poisson model: 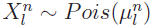 for *l* = 1, …, *L*. We allowed the mean read profile 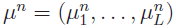 to vary across sites by using a hierarchical version of the multi-scale model for inhomogeneous Poisson processes introduced by (Kolaczyk, 1999) and (Timmermann and Nowak, 1999).

To introduce the ideas behind the multi-scale model, consider a single site with parameter vector *μ* = (*μ*_1_, …, *μ*_*L*_) (so drop the superscript ^*n*^ for simplicity). The key idea behind multi-scale Poisson models is to reparameterize this model in terms of parameters that capture spatial variation in *μ* at multiple scales, as follows. Let 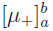 denote the sum 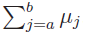. At the “zeroth” scale, define a single intensity parameter *λ*_0_ that captures the total intensity in the region

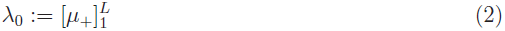

At the first scale define a single parameter that captures the relative intensity in the first half of the region vs the entire region:

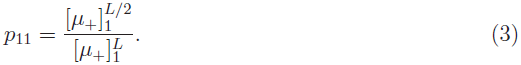

At the second scale, define two parameters: one that captures the relative intensity in the first quarter of the region vs the first half; and one that captures the relative intensity in the third quarter vs the second half.

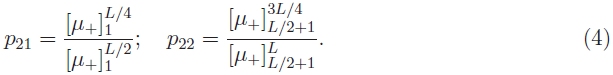

At the third scale there are four parameters *p*_31_, …, *p*_34_ that similarly capture the relative intensity of an eighth of the region vs each quarter. This continues up to the *J*th scale (where recall *J* = log_2_(*L*)), in which there are *L*/2 = 2^*J*−1^ parameters of the form

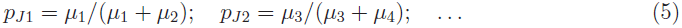

Combining across scales 0 to *J* this defines a total of *L* parameters, *p* = (*λ*_0_, *p*_11_, *p*_21_, *p*_22_,…., *p*_*J*(*L*/2)_), which are a one-one function of *μ*. That is, this defines a reparameterization of the model from *μ* = (*μ*_1_, …, *μ*_L_) to *p* = (*λ*_0_, *p*_11_, *P*_21_, *P*_22_, …, *P*_*J*(*L*/2)_).

This reparameterization has two key features: i) the likelihood P(*X|p*) factorizes into a product form over the *L* elements of *p* (just as the likelihood P(X|*μ*) factorizes into a product over the *L* elements of *μ*). Indeed, from elementary properties of the Poisson distribution, this factorization includes a Poisson likelihood for *λ*_0_ and a Binomial likelihood for each of the other parameters in *p*; see Supplementary Methods and Kolaczyk (1999) for details. ii) spatially-structured perturbations to the vector *μ* are captured by large perturbations in just a few elements of *p*. (By a spatially-structured perturbation, we mean a modification *μ*_*i*_ → *μ*_*i*_ + *δ*_*i*_ such that *δ*_*i*_ tends to be similar to *δ*_*j*_ when |*i*—*j*| is small.) This property is related to the similar key property of wavelets (Donoho and Johnstone, 1995), which are perhaps the best known multi-scale methods: spatially smooth signals tend to be concentrated into a small number of wavelet coefficients.

As a consequence of ii) we modeled spatially-smooth heterogeneity in *μ*^1^, …, *μ*^*n*^ across putative binding sites using a simple hierarchical model for *p*^1^, …, *p*^*n*^ (where we have reintroduced superscript ^*n*^ to index sites). Specifically, we introduced parameters 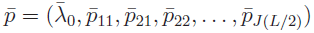 to represent the mean cleavage pattern across sites, and then assumed that site specific parameters *p*^1^, … , *p*^*n*^ are independent and identically distributed given 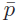, with

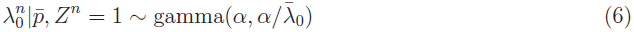

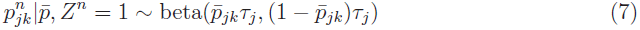

for *k* = 1, …, 2^*j*−1^ and *j* = 1, …, *J*, where *α* and *τ*_*j*_ are hyperparameters that control variability in the parameters at different scales.

### 2.2 msCentipede model at unbound motifs

We modeled the read count profile at the *n*^th^ site *X*^*n*^ conditional on *Z*^*n*^ = 0 using the same Poisson model, but with different distributions for the parameters:

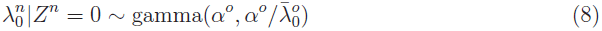

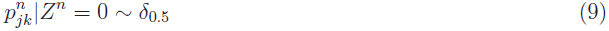

where *δ*_0.5_ denotes the distribution with point mass on 0.5. Note that this means that 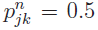, which is equivalent to assuming that the Poisson rates *μ* = (*μ*_1_, …, *μ*_*L*_) are all equal, resulting in uniformly distributed reads over the entire site. That is, it corresponds to the commonly-used assumption that there is no spatial structure in the read count profile when the transcription factor is not bound to its motif. (See Section 2.4 for a more flexible model for unbound sites.)

### 2.3 CENTIPEDE is a special case of msCentipede

The above msCentipede model (8–9) for unbound sites is exactly the same as the CENTIPEDE model for unbound sites. (The assumption of a gamma distribution for the Poisson rate parameter 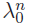 in (8) implies a negative binomial distribution for the total read-counts, which is exactly the model assumed by CENTIPEDE.) Furthermore, the msCentipede model for bound sites (7) becomes equivalent to the original CENTIPEDE model for bound sites in the special case *τ*_*j*_ → ∞, which corresponds to no heterogeneity in the shape of the cleavage pattern across bound sites. That is, msCentipede is an extension of CENTIPEDE to allow for heterogeneity in the shape of the cleavage pattern across bound sites.

### 2.4 Flexible model for background DNase I cleavage rate

A number of studies have highlighted a strong sequence preference for DNase I cleavage (Hesselberth *et al.,* 2009; Koohy *et al., 2013;* He *et al.,* 2014). This sequence preference would cause the distribution of reads at unbound motif instances to be i) systematically non-uniform near the shared core motif; and ii) varying among motif instances away from the shared core motif (due to differences in the surrounding sequence). To account for these factors we consider a more flexible model for unbound sites. Specifically, we modify (9) as follows:

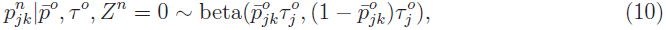

where the background parameters 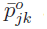 and τ^*o*^ control the mean profile and the variance about this mean respectively. We estimated these background parameters using DNase-seq reads from naked DNA around the same set of motif instances, and refer to the method using this more flexible background model as msCentipede-flexbg. (In principle it is also possible to estimate these parameters using the DNase-seq data from chromatin, as part of the clustering of motif instances into bound and unbound motifs, but when we tried this we found msCentipede performed worse in practice than the uniform model (9), presumably because of the cost associated with attempting to estimate the many additional parameters of this more flexible model; see Discussion).

### 2.5 msCentipede when multiple replicates are available

When multiple replicates are available, msCentipede treats the replicates as independent samples. Ideally, it is desirable to model the heterogeneity across genomic sites and the heterogeneity across replicates separately. However, in practice, we usually have only two or three replicate DNase-seq measurements in a given cell type, making it difficult to accurately quantify the heterogeneity across replicates. Instead, our approach assumed that heterogeneity across replicates and heterogeneity across genomic sites were the same (see Supplementary Methods for more details).

### 2.6 Parameter estimation and inference

The parameters in msCentipede, *ζ*, *α*, 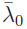, *α*^*o*^,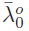, *τ*_*j*_,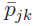, *k* = 1, …,2^*j*−1^ and *j* = 1, …, *J*, were estimated by maximizing the likelihood across all putative binding sites using an expectation-maximization algorithm (see Supplementary Material for details). When DNase-seq data assayed in naked DNA were available, the background parameters 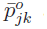 and 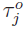 were first estimated using naked DNA assays; keeping these fixed, we, then, learned the remaining model parameters.

Inference on binding sites can be performed by computing the posterior odds for each site:

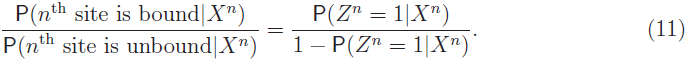

Detailed computation of P (*Z*^*n*^ = 1|*X*^*n*^) is given in Supplementary Methods.

## 3 Applications

In this section, we evaluate the accuracy of msCentipede on a set of transcription factors for which high quality ChIP-seq data and highly informative position-weight matrix (PWM) models are available. We also evaluate the gain in performance achieved when we use a more flexible model for background DNase I cleavage rate, with parameters for this model learned using DNase-seq data from naked DNA.

### 3.1 Description of data and validation metrics

We executed msCentipede and CENTIPEDE using DNase-seq and ATAC-seq measurements assayed in the GM12878 lymphoblastoid cell line as data. Two replicate measurements using the UW DNase-seq protocol (Hesselberth *et al.,* 2009) and four replicate ATAC-seq measurements (Buenrostro *et al.,* 2013) were available for this cell line. The DNase-seq data were single-end reads that can be converted to counts of DNase I nicks for each base pair in a straightforward manner. The ATAC-seq data were paired-end reads; however, we ignored the information in the length of DNA fragments and used the counts of transpositions for each base pair as data. In addition, we executed PIQ (Sherwood *et al.,* 2014) using DNase-seq data to compare its performance with msCentipede.

We compared the three algorithms on a set of 50 transcription factors with ChIP-seq data assayed by ENCODE in the same cell line (ENCODE, 2012), and for which PWM models were computed using data from high-throughput SELEX experiments (Jolma *et al.,*2010, 2013). For each transcription factor, we identified a genomewide set of high-quality putative binding sites (PBS) using human genome reference GrCh37; for each PBS, the likelihood ratio for the PWM model vs a background model exceeded 1000. Using a 64 base-pair window around each PBS, we filtered out those sites that had fewer than 80% of bases in their window to be uniquely mappable. For each of the remaining sites, we computed the posterior probability that the transcription factor is bound, using CENTIPEDE and msCentipede. We used DNase-seq read count data from naked DNA derived from the IMR90 cell line (Lazarovici *et al.,* 2013) to fit the background model parameters in msCentipede-flexbg. In the case of PIQ, we used the “score” output by the algorithm as a measure of confidence of whether a motif instance is bound. When multiple replicate measurements are available, we executed PIQ by providing data from the replicates as separate input files.

We evaluated the accuracy of each algorithm using Area under the Receiver Operating Curve (AuROC). To compute the AuROC, we selected a gold standard set of ‘bound motif instances’ and ‘unbound motif instances’; bound motif instances were PBS that lied within a ChIP-seq peak identified by a peak caller and ‘unbound motif instances’ were PBS that lied outside ChIP-seq peaks and had fewer ChIP reads than reads from a control IP experiment mapping to a 400 base pair window around the PBS, after controlling for total read depth. For each transcription factor, we executed two peak callers, MACS (Zhang *et al.,* 2008) and GEM (Guo *et al.,* 2012), each with a 1% FDR cutoff, to generate two gold standard sets of bound and unbound motif instances. In this paper, we illustrate the accuracy of the algorithms evaluated against gold standards generated using MACS when using DNase-seq measurements as data. The accuracy of all three algorithms improved by a modest amount when using the gold standards generated by GEM, presumably due to the use of sequence information by GEM to better identify ChIP-seq peaks (see Figure S2).

### 3.2 Results

#### 3.2.1 msCentipede achieves improved accuracy

msCentipede achieved AuROC comparable to or better than CENTIPEDE across a broad range of transcription factors when each algorithm was applied to chromatin accessibility measurements from a single DNase-seq assay as shown in Figure 2a. Compared with PIQ, we observed that msCentipede achieved substantially higher AuROC for some factors and lower AuROC for others, as shown in Figure 2c. When multiple replicates are available, CENTIPEDE treats them by pooling the replicate datasets; however, msCentipede treats replicates by modeling them as independent samples. By modeling the replicates appropriately and accounting for heterogeneity across genomic sites and replicates, msCentipede achieved substantial increase in AuROC compared to CENTIPEDE and PIQ for a broad range of transcription factors, as illustrated in Figures 2b and 2d. Similar improvements in accuracy for msCentipede compared to CENTIPEDE were observed when using ATAC-seq measurements as data (see Figure S3).

**Figure 2:**
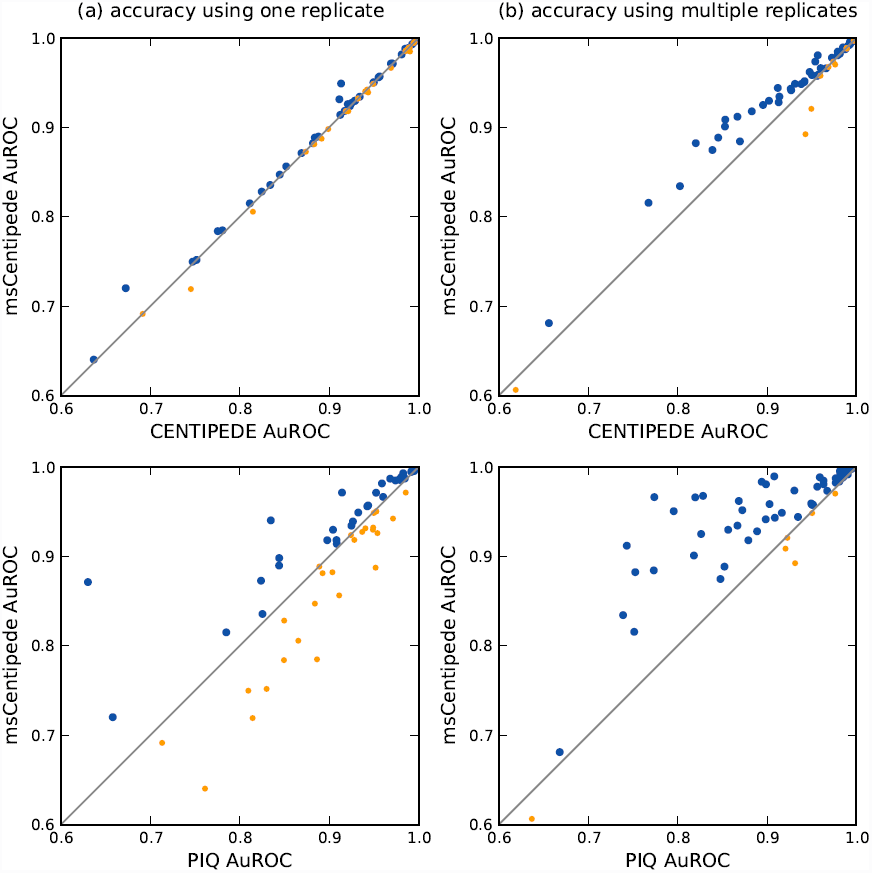
Accuracy of msCentipede, CENTIPEDE and PIQ across a range of transcription factors. Each point corresponds to a different factor and accuracy is measured by area under the ROC curve. Blue points correpond to factors where msCentipede achieves higher accuracy than CENTIPEDE (top panels) or PIQ (bottom panels), and orange points correspond to a worse performance by msCentipede. In the left panels (a), the algorithms are compared using data from a single replicate, while the right panels (b) show results when data from multiple library replicates are analyzed.

For each transcription factor, the hyperparameter *τ* gives a measure of heterogeneity in read distribution across genomic sites and replicates, with lower values indicating greater heterogeneity. In Figure S4, we observed that the values of the hyperparameters *τ*_*j*_ were rather small, suggesting that we were able to increase power by better modeling variation in the data. Furthermore, as expected, we observed a higher degree of overdispersion in read distribution at finer resolutions compared to coarser resolutions for a number of transcription factors.

#### 3.2.2 Modeling DNase I cleavage patterns improves factor binding inference

In recent work, He *et al.* (2014) and Sung *et al.* (2014) demonstrated that strong DNA sequence preference for DNase I cleavage could pose a challenge to using the detailed shape of DNase cleavage profiles for inferring transcription factor binding. Specifically, He *et al.*(2014) identified motif instances that lie within peaks in ChIP-seq measurements for a transcription factor in a given cell line. Using these instances, they showed that, in a region of ~20 bp surrounding the motif, the mean DNase I cleavage profile estimated from naked DNA (unbound sites) matched the mean cleavage profile estimated using DNase-seq data from the same cell line (bound sites). Starting from similar observations, Sung *et al.* (2014) clarified that although sequence-preference effects were evident for all transcription factors, some transcription factors - those with slower-binding kinetics - show an appreciable reduction in the cut profile around the bound motifs (a “footprint”), whereas others - those with faster-binding kinetics - show little or no footprint.

These observations raise two questions: first, whether the uniform background model (assumed by CENTIPEDE, and msCentipede) for the unbound sites might be better replaced by a non-uniform background model capable of capturing the sequence preference effects around the motifs; second, whether it might be better to entirely ignore the DNase I cleavage profile when attempting to distinguish between bound and unbound sites - and, rather, to focus only on the total intensity of DNase I hypersensitivity in the region. To test this, we compared the accuracy of three different models for transcription factor binding:

1. ‘no cleavage profile’ model that ignores the cleavage profile, and simply models the total DNase read counts using Poisson-gamma distributions at bound and unbound sites (described earlier).
2. msCentipede
3. msCentipede-flexbg, which allows for a non-uniform background model, with parameters estimated using DNase-seq measurements from naked DNA around the same set of PBS.

Comparing first the msCentipede model with the no-cleavage model, we found the accuracy of msCentipede to be substantially greater for a broad range of transcription factors (Figure 3a). This result may appear to conflict with results from He *et al.* (2014); Sung *et al.*(2014) showing that cleavage patterns within factor-bound motif instances are driven primarily by sequence preferences for DNase I cleavage, which suggests that use of the cleavage profile to identify binding sites could increase false positive findings. However, we note that i) sequence preference effects, while presumably occuring genome wide, are *shared* across binding sites only in the small region around the shared sequence motif (typically 10 − 20 bp), while most methods to detect factor binding, including ours, make use of cleavage patterns in much larger windows (typically 50 − 100 bp) around the motif instance, and ii) for some factors – those with slower binding kinetics – the footprint effect (i.e. the systematic overall decrease in DNase signal surrounding the motif) may be helpful in distinguishing bound and unbound sites, and the benefits of this could outweigh the unmodelled sequence preference effects.

**Figure 3:**
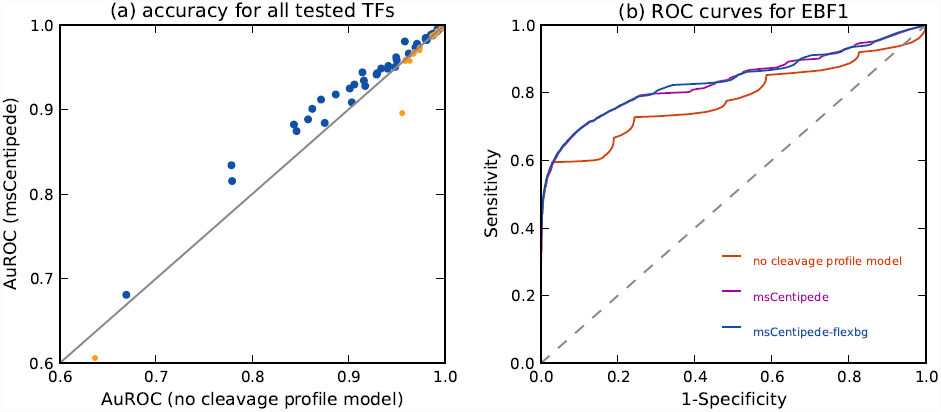
Modeling factor-specific DNase I cleavage profile and sequence bias in DNase cleavage increases prediction accuracy. (a) Modeling the DNase I cleavage profile at bound sites increases the prediction accuracy of msCentipede across a broad range of transcription factors. Each point on the plot corresponds to a different transcription factor. (b) We show the ROC curves for transcription factor EBF1 for three different models of increasing complexity. We observe a substantial increase in accuracy when incorporating a multi-scale model for the factor-specific cleavage profile; however, the increase in accuracy when modeling the background cleavage rate using naked DNA data is rather modest. This holds true for a broad range of factors as shown in supplementary Figure S6.

We turn now to the comparison of msCentipede with msCentipede-flexbg. Note that msCentipede-flexbg, by modeling the background cleavage profile using naked DNA assays, has the potential to eliminate false positives due to sequence-driven cleavage patterns highlighted by He *et al.* (2014); Sung *et al.* (2014). And indeed, we found that, for most factors, the estimated mean background cleavage profile, captured by the parameters 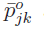, was non-uniform within the motif, reflecting precisely the sequence preferences for DNase I cleavage (Figure S5). However, we also found that this improved background model resulted in only modest improvements in accuracy of identifying bound sites (Figure S6).

Using transcription factor EBF1 as an example, Figure 3b illustrates that all three models have very similar true positive rates up to a false positive rate of 3 − 4%. However, incorporating the DNase cleavage profile substantially increased the true positive rate for false positive rates larger than 4%. This suggests that while modeling the total DNase read counts alone was sufficient to accurately identify bound PBS with highest total DNase-seq signal, incorporation of the DNase cleavage profile was necessary to identify bound PBS with moderate total DNase-seq signal. These PBS may be indicative of low occupancy sites where the binding of the transcription factor is in a less stable equilibrium and the factor is likely bound to the DNA at these PBS in a smaller fraction of the cells assayed.

## 4 Discussion

We developed msCentipede, a hierarchical multi-scale model to accurately identify binding of a transcription factor using sequencing reads from DNase-seq or ATAC-seq assays and the sequence content of putative binding sites for that factor in the genome. While previous approaches like CENTIPEDE have successfully used the characteristic profile of DNA hypersensitivity to DNase I around bound motif instances to identify factor binding sites, the multinomial model used in CENTIPEDE ignores spatial structure in the data and makes a strong assumption on the heterogeneity in read distribution across bound sites in the genome. The hierarchical multi-scale model explicitly allows for heterogeneity in the read distribution across bound sites (with different amounts at different scales), resulting in a substantial increase in accuracy across a broad range of transcription factors. Finally, we proposed a more flexible background model that requires the availability of DNase-seq (or ATAC-seq) data assayed in naked DNA. This flexible background model has the potential to account for heterogeneity in background DNase I cleavage rate specific to the sequence context of motif instances of the transcription factor.

A simple extension to CENTIPEDE that can account for heterogeneity across sites is to allow for site-specific parameters in the multinomial distribution and to model these site-specific parameters using a Dirichlet distribution. However, this multinomial-Dirichlet model is not sufficiently flexible to capture potential spatial structure in heterogeneity in DNase I cleavage. The proposed multi-scale model allows different amounts of heterogeneity across different scales and effectively captures spatial structure in the heterogeneity. It is fairly straightforward to extend the proposed framework to model spatial structure in the mean cleavage pattern 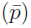 as usually modeled in multi-scale approaches (Kolaczyk, 1999; Donoho and Johnstone, 1995; Shim *et al.,* 2014). However, we found that this extension was computationally expensive and gave very minor improvements in accuracy, presumably because there were so many motif instances that we could accurately estimate the mean pattern without spatial smoothing.

When considering a flexible model at unbound motif instances which allows for spatial structure and heterogeneity in background DNase I cleavage patterns, it is natural to estimate the parameters of this model using data from the relevant cell type. However, we observed that when all the parameters in the flexible model are estimated using data from chromatin, the model tended to estimate smaller values for the precision parameter, *τ*, at bound sites resulting in a large number of ‘true’ unbound sites being incorrectly identified as bound.Currently, we suggest using the flexible model (msCentipede-flexbg) only when DNase-seq (or ATAC-seq) data assayed in naked DNA is available. However, a framework that allows estimation of heterogeneity in background DNase I cleavage from data assayed in the relevant cell type may be be more accurate.

msCentipede-flexbg estimates spatial structure and heterogeneity in the background model using DNase-seq data from naked DNA at all motif instances; thus, the heterogeneity in background read distribution is primarily driven by variation in sequence context around motif instances. However, within a cell type, variation in background chromatin context at unbound sites (e.g., whether the motif instance is in the linker region or in DNA wrapped around a nucleosome, and which other transcription factors are bound at or close to the motif instance) is likely to be a larger source of heterogeneity in background read distribution than variation in sequence context. This intuition suggests that we should estimate the precision parameter at unbound sites *τ*^*o*^ using DNase-seq data from chromatin, rather than using DNase-seq data from naked DNA. However, using this approach, we observed the background precision parameter *τ*^*o*^ in msCentipede-flexbg was consistently underestimated when this parameter was estimated using data from chromatin, resulting in a high false positive rate. Extensions to these models that accurately capture the background heterogeneity in the data across genomic sites would be a useful avenue for future research.

